# Bursting out: linking changes in nano-topography and biomechanical properties of biofilm-forming Escherichia coli to T4 lytic cycle

**DOI:** 10.1101/2020.06.29.176883

**Authors:** Shiju Abraham, Yair Kaufman, François Perreault, Ry Young, Edo Bar-Zeev

## Abstract

Bacteriophage infection cycle has been extensively studied, yet little is known on the structural and mechanical changes that lead to bacterial lysis. Here, bio-atomic force microscopy was used to study in real-time and *in-situ* the impact of the canonical phage T4 on the nano-topography and biomechanics of irreversibly attached, biofilm-forming *E. coli* cells. The results show that in contrast to the lytic cycle in planktonic cells, which ends explosively, anchored cells that are in the process of forming biofilms undergo gradual lysis, developing distinct sub-micron lesions (∼300 nm in diameter) within the cell envelope. Furthermore, it is shown that the envelope rigidity and cell elasticity decrease (>50% and >40%, respectively) following T4 infection. These new insights show that the well-established lytic pathway of planktonic cells may be significantly different from that of biofilm-forming cells. Elucidating the lysis paradigm of these cells may advance biofilm removal and phage therapeutic.

There is no conflict of interest and all co-authors have seen and approved the current version for submission.

## INTRODUCTION

Caudovirales are dsDNA tailed phages that dominate the virosphere and are being developed as alternative therapeutics against multi-drug resistant bacterial infections. The lysis event that terminates the Caudovirales infection cycle has been studied primarily in model phages of planktonic *E. coli* cultures, especially lambda and T4. The *E. coli* host is a Gram-negative bacterium, comprising a ∼ 45 nm thick cell envelope that includes a peptidoglycan layer sandwiched between inner and outer membranes ^1^. *E. coli* bacteria are ubiquitous in the environment and, in the planktonic state, these cells are usually harmless to humans. However, some *E. coli* strains have acquired the ability to form pathogenic biofilms that cause a broad spectrum of diseases ^2,3^. In the initial stages of *E. coli* biofilm formation, attached cells secrete extracellular polymeric substances (EPS) that act as a sticky scaffold for anchoring to the conditioned surface and to each other ^3,4^. Once developed, *E. coli* biofilms are notoriously resistant to removal by antibiotics due to the protective barrier provided by the EPS matrix ^3^.

An alternative to antibiotic treatment for *E. coli* biofilms is the use of bacteriophage, especially since many coliphages are equipped with virion-mounted enzymes that can degrade elements of the EPS matrix ^5,6^. However, details of the phage lytic cycle in biofilm state are lacking, especially in terms of host lysis and virion dissemination. Microscopic real-time imaging has indicated that in the planktonic state, infected *E. coli* cells lyse explosively, usually from a single point on the rod-shaped cellular envelope ^7^. Results from biochemical studies, genetics and fluorescence microscopy have led to a three-step model for the lysis pathway, in which different phage-encoded proteins called holins, endolysins, and spanins sequentially target each of the three layers of the *E. coli* envelope ^8^. The final step in this lytic cycle is localized fusion of the inner and outer membrane after the temporally regulated degradation of the peptidoglycan layer. This last step in the lytic cycle leads to the catastrophic explosion of the planktonic cell.

Although the lysis event has been well-studied operationally through the use of video-microscopic as well as molecular and physiological methods ^9–11^, there is little information on the impact of the lytic pathway on biomechanics and physical structure of the infected cell. During the last decade, several researchers have used atomic force microscopes (AFM) to characterize the infection cycle of *E. coli* by different phages. Initially, AFM imaging of infected and dehydrated *E. coli* revealed that by ∼30 min after T4-phage infection, the cell envelope has undergone various topographical changes ^12^. Later, AFM imaging in more physiologically relevant conditions was done for cells infected with the non-lytic filamentous phage M13, where the progeny are extruded through the intact envelope ^13^. Although no change in cell morphology was detected, force vs. indentation measurements have shown that the Young (elastic) modulus of these *E. coli* cells decreased by ∼57% after M13 phage infection ^13^. For lytic infections, quantitative and kinetic information over the mechanical properties of bacteria is lacking despite the centrality of this process in phage therapeutics. Moreover, most of the molecular and cellular studies to date have been done with planktonic cells, and there are reports indicating that the phage infection pathway may be significantly different in biofilms ^14,15^.

Here, we characterized the nano-topography and analyzed the biomechanical properties of *E. coli* cells that were infected by T4 phages while in the process of forming a biofilm. Using bio-AFM, we acquired nano-topography images of the *E. coli* envelope as well as force vs. indentation curves of the entire cell in real-time under physiological conditions. These measurements allowed us to calculate the elastic modulus of *E. coli* cells before, during, and after T4 infection. Our results provide a direct link between changes in the envelope and cell structure to biomechanical properties of *E. coli* cells during the lytic cycle of T4 phages.

## RESULTS

### Changes in nano-topography of biofilm forming *E. coli* cells during T4 phage infection

*E. coli* culture was grown on a glass AFM coupon that was pre-coated with a ∼5 nm thin, rigid ^16^ and positively charged LBL (Fig. 1). Over few hours (8 h), *E. coli* cells were irreversibly anchored to the AFM coupon, with some having developed into monolayer clusters, which are early stages in biofilm formation ^17^. Irreversible cell attachment to the LBL glass coupon enabled us to capture real time AFM images and acquire force measurements in a physiological solution. In contrast, cells applied without the LBL were sparsely attached and dislodged from the surface by the AFM tip, thus hindering images acquisition and force measurements. Live-dead staining experiments indicated that attachment to the LBL had no impact on *E. coli* viability (Fig S1).

**Figure 1.**
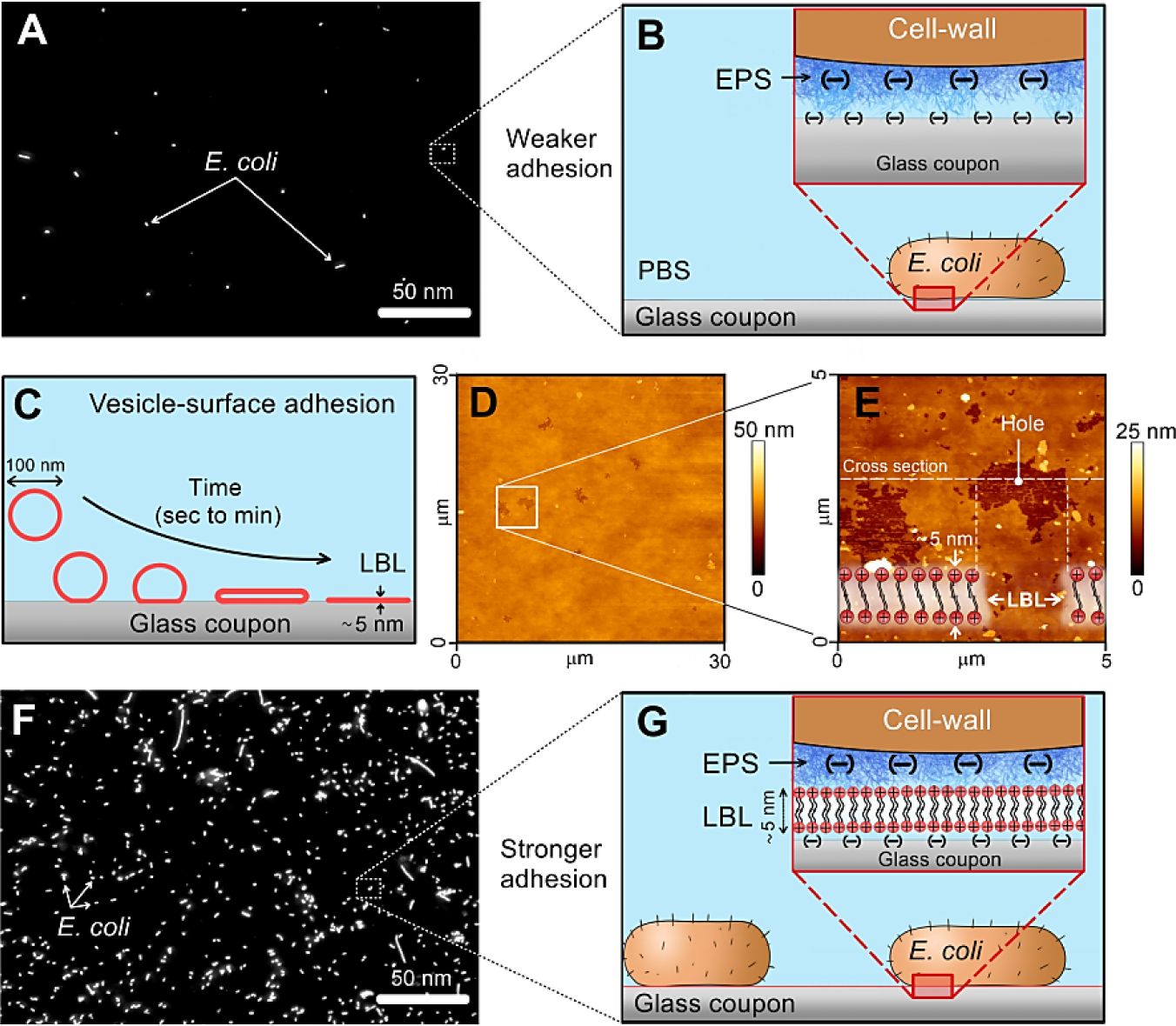
*E. coli* cells were attached to an AFM glass coupon via a positively charged lipid bilayer (LBL), which served as a rigid and adhesive support. (A) Fluorescence micrograph of a sparse *E. coli* cell coverage on a glass coupon without LBL coating. (B) Schematic illustration of the molecular interactions between *E. coli* cells and the glass coupon, pointing on repulsive electrostatic interactions together with weak van der Waals adhesion between the extracellular polysaccharide substances (EPS) and the surface. (C) LBL was prepared on an AFM glass coupon using vesicle fusion techniques, where lipid vesicles adhere to the surface, rupture, and form an LBL coating. (D, E) AFM images (in physiological solution) of positively charged LBL on an AFM glass coupon with a corresponding illustration of the LBL coating. Holes in the LBL allowed us to measure the layer thickness. (F) Fluorescence images of *E. coli* cells attached to a glass coupon that was pre-coated by a positively charged LBL. (G) Schematic illustrations that exhibit the electrostatic and weak van der Waals adhesion that attaches *E. coli* cells to the positively charged coated surface.

The topography and viability of irreversibly attached *E. coli* cells before and after T4 infection were studied *in-situ* by epifluorescence microscopy as well as by AFM and TEM (Figs. 2, 3). Initially, bacteria were fluorescently tagged by a Live/Dead staining kit, namely with SYTO9 (green) and PI (red). From this initial stage and up to 10 min after the addition of T4 phages, all the cells exhibit green (“live”) fluorescence, indicating that the cell membranes were intact (Fig. 2A). Three dimensional AFM image analysis of uninfected bacteria indicated that cell size did not change in comparison to their dimensions prior to T4 addition; with an average width of ∼1 µm and a length that ranged from 2 µm to 3 µm (Fig. S2). We note that attached phages were not detected by AFM under these *in-situ* conditions, probably because the AFM tip has physically removed the T4 from the surface of *E. coli* cells during the scan. Nonetheless, TEM images from concomitant subsamples indicated that one to two phages were attached per cell by 10 to 20 min after the addition of the phages (Fig. 2A, right panel).

**Figure 2.**
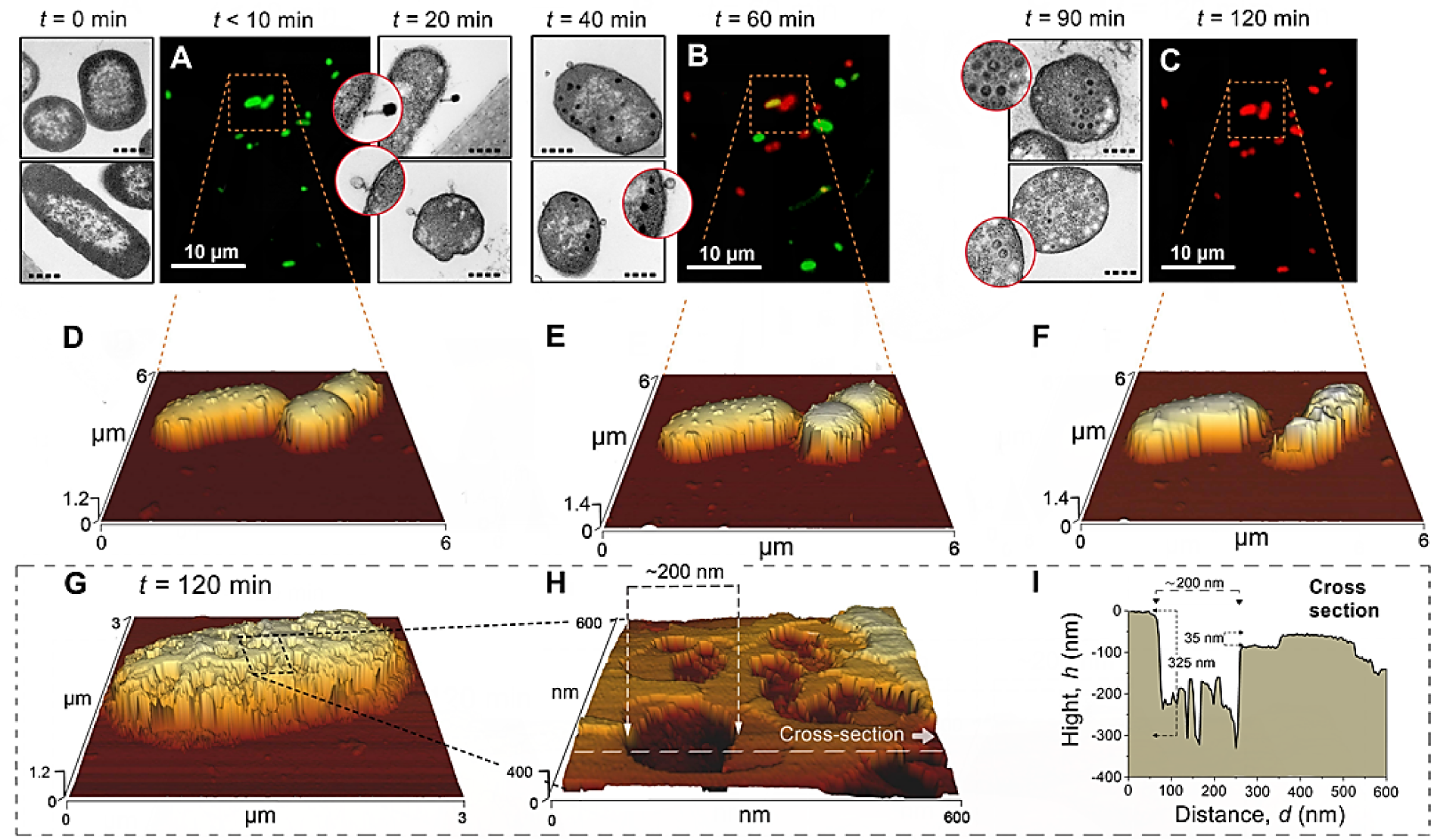
(A-C) *In-situ* fluorescence and (D-H) concomitant AFM images following the infection of biofilm forming *E. coli* cells by T4 phages. (I) Cross-sectional analysis of the corresponding AFM nano-topography image (H). Epifluorescence images of “live” (green) and “dead” (red) cells, as well as corresponding AFM scans, which were captured immediately after T4 infection (*t*<10 min) as well as 60 min, and 120 min following T4 addition. “Live” and “dead” bacteria were identified by staining the cells with SYTO 9 and PI, respectively (see also Fig. S4). Complementary TEM images were captured before T4 addition as well as 10, 20, 40 and 90 min after T4 were added. Black circles within the TEM inserts (*t*= 40, 90 min) indicate the presence of T4 virions inside the cells. TEM scale bar was 1 µm (full-line) for the low magnification and 0.4 µm (dash line) for the larger magnifications. Similar images were captured from 12 other cells at 5 individual experiments.

**Figure 3.**
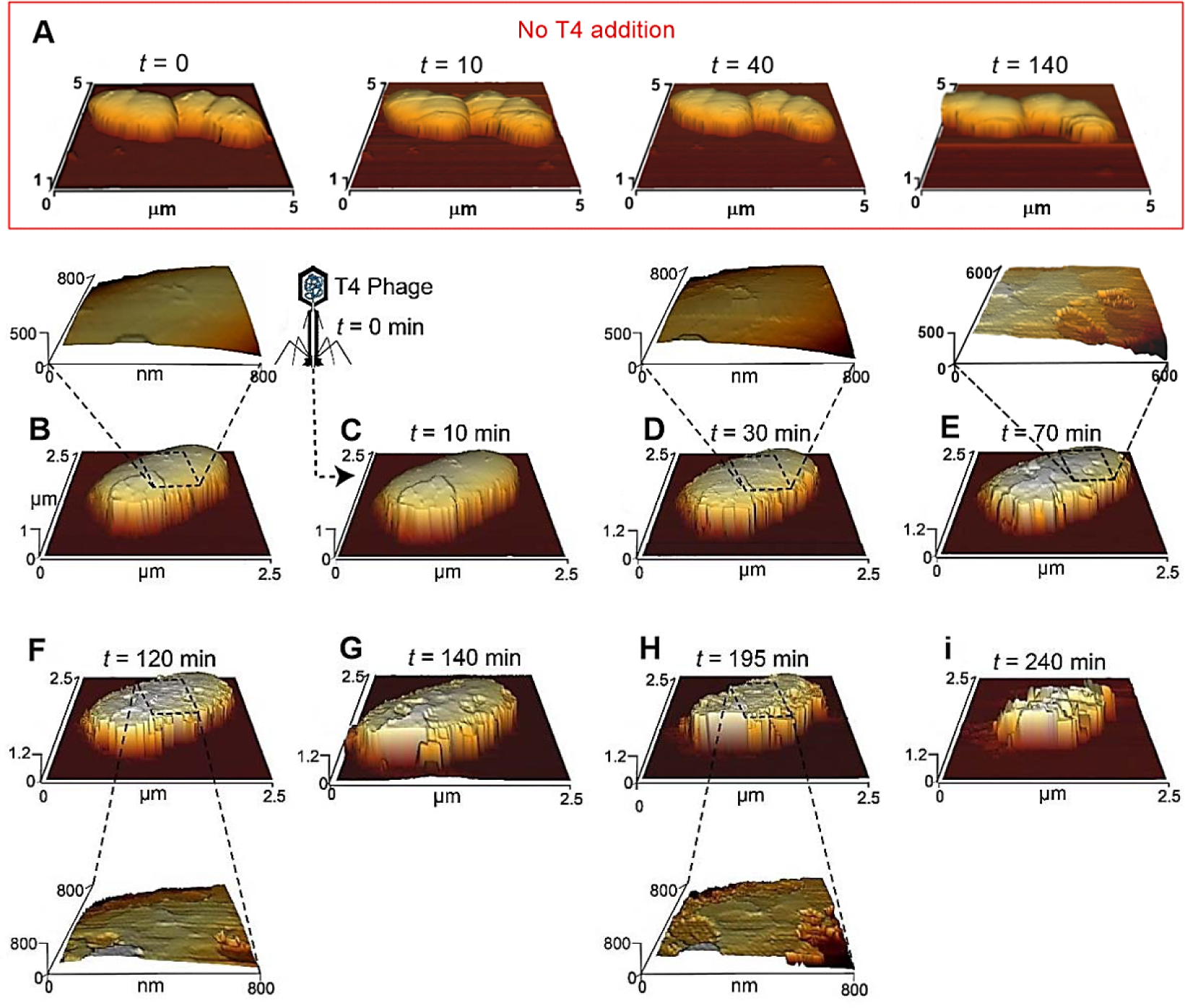
Three dimensional images of an *E. coli* cell during T4 phage infection captured *in-situ* by AFM. (A) Time series images of uninfected *E. coli* cells, namely with no addition of T4. (B) The first image was captured before the addition of T4 phages. (C-i) Time series of AFM images that show the changes in nano-topography of the same cell during the infection cycle under hydrated conditions. Similar images were captured from 12 other cells at 5 individual experiments.

By one hour after the addition of T4 phages, both “live” (green) and “dead” (red) cells were captured, indicating that the cytoplasmic membrane of the latter has been permeabilized (Fig. 2B). In some cells, yellow fluorescence was also detected, presumably reflecting that the membrane has been permeabilized but the PI stain has not yet completely reduced the SYTO 9. The fluorescence intensity of cells that were stained by SYTO 9 increased with time; 50% ± 25% brighter at *t*=60 min compared to *t*=0 (Fig. 2A, B), indicating that phage DNA replication was robust. Complement AFM nano-topography images indicated that *E. coli* were mostly intact with no significant change to their size (Fig. 2E). Yet, during this stages (40-90 min), up to 20 virions were detected within 20-60% of the cells captured by TEM (Fig.2B-C, left panels).

Nonetheless, two hours after T4 addition, most of the cells were stained with PI (red, Fig. 2C) or completely demolished (Fig. 2F). Concomitantly, AFM images indicated that *E. coli* cells were structurally deformed compared to the same cells at *t*=60 min (Fig. 2F). Moreover, the nano-topography of infected bacteria at *t*=120 min shows that cell envelopes were perforated with distinct lesion of 100-300 nm and up to ∼350 nm in depth (Fig. 2G-I). Often, the lesions/ruptures had shallow ledges (few tens of nm) with much deeper crevices that may reach a few hundreds of nm (Fig. 2 H-I). It should be noted, according to AFM images of un-infected *E. coli* bacteria, no damage to the cell envelope was caused by the AFM tip during imaging (Fig 3A).

Dynamic, changes in the nano-topography of the cell structure during T4 phage infection were observed *in-situ* by sequential AFM images (Fig. 3). Uninfected cells remained similar with no changes to the nano-topography of the cell envelope or growth over two hours (Fig. 3A). In contrast, during the first 10 to 30 min after the addition of T4 (Fig 3C, D), the definition of the envelope topography was evident compared to uninfected cells (Fig. 3A), as if the cell capsule was removed. Yet, no apparent changes to envelope nano-topography were captured (Fig 3C). However, 70 min after the addition of T4, some superficial ruptures (diameter of 105-190 nm and 10-15nm depth) were visualized in the cell envelope (Fig. 3E). More cavities accumulated over the course of the infection (Fig. 3F-H), ultimately culminating in cells with irregular boundaries and dimensions less than half of the original cell (Fig. 3i).

### Biomechanical changes of *E. coli* cells during T4 phage infection

Changes in the biomechanics of the cell envelope (Fig. 4) and the entire bacterium (Fig. 5) following T4 infection were measured *in-situ* by bio-AFM. The stiffness of the cell envelope was calculated as Δ*F/*δ, where *F* is the force that was applied on the cell and δ is the indentation of the AFM probe into the cell. Measurements were averaged from a total of ∼45,000 data points taken at 4 to 5 different locations (250-500 nm^2^) on the surface of each bacterium (*n*=12) during infection (Fig. 4 A,B). These force measurements were done using a pyramid AFM tip with a radius of curvature of ∼5 nm, and the approach speed was 190±20 µm/s. A significant reduction in cell envelope stiffness (16%, *p* < 0.02) was already detectable 10 to 20 min after the addition of T4 phages and continued to decrease monotonically, reaching ∼58% reduction by two hours (Fig. 4D). In contrast, no change in cell envelope stiffness was detected over a similar time frame for uninfected bacteria (Fig. 4D).

**Figure 4.**
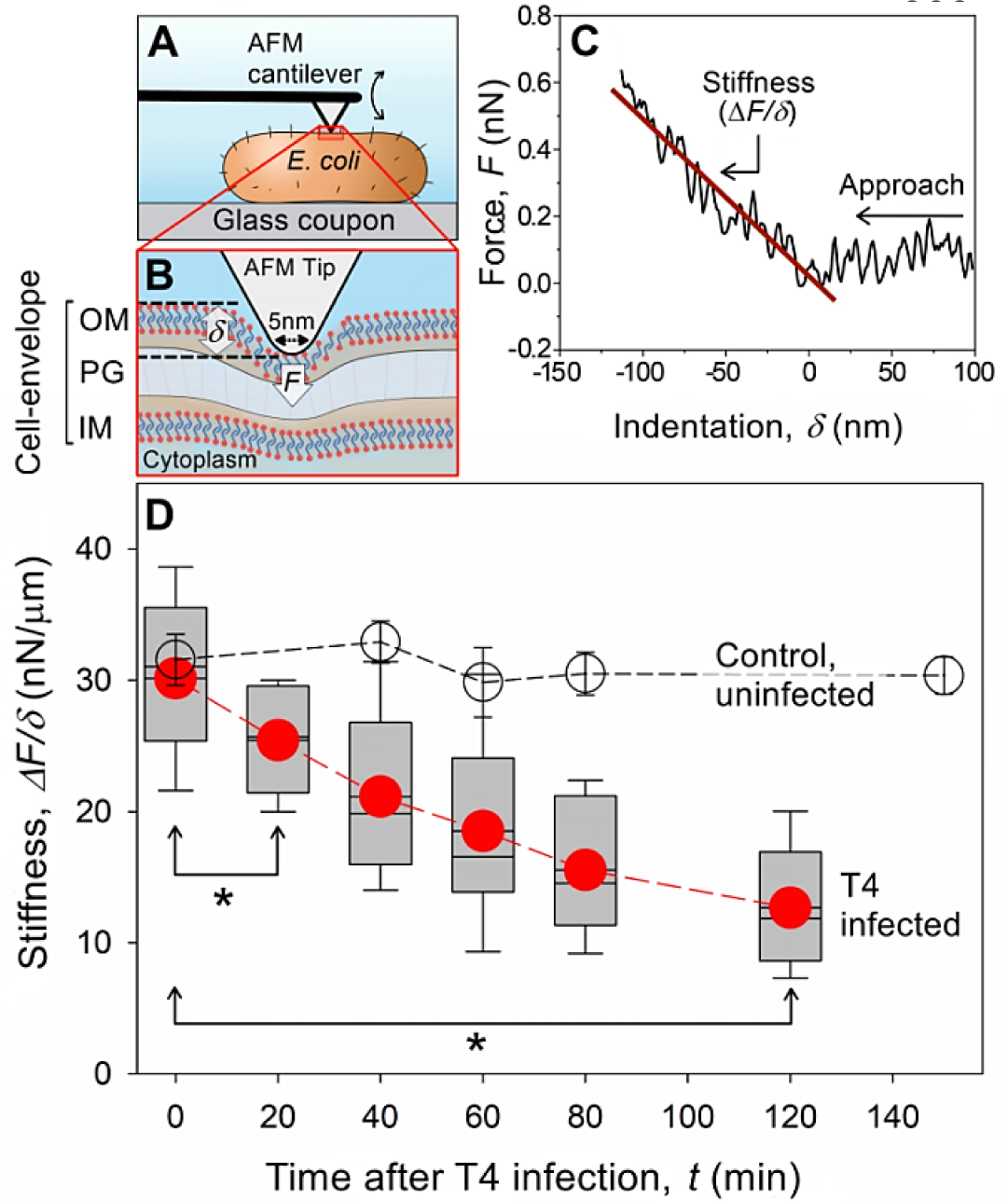
Stiffness measurements of *E. coli* cells during T4 infection. (A and B) schematic illustration of AFM stiffness measurement using a pyramid AFM tip (not to scale). (C) A representative AFM force vs. indentation that demonstrates how the stiffness was calculated. Approach velocity was 190± 20 µm/s (D) Stiffness (rigidity) changes with time for different bacteria with (box plot with red circles indicating the average) and without (empty circles) the addition of T4 phages. Average stiffness was based on 45,000 force measurements taken from 4 to 5 areas on the surface of each bacterium. Asterisks indicate a statistically significant difference between two time points (*p* < 0.01, *n*=12 at 3 to 5 independent experiments).

**Figure 5.**
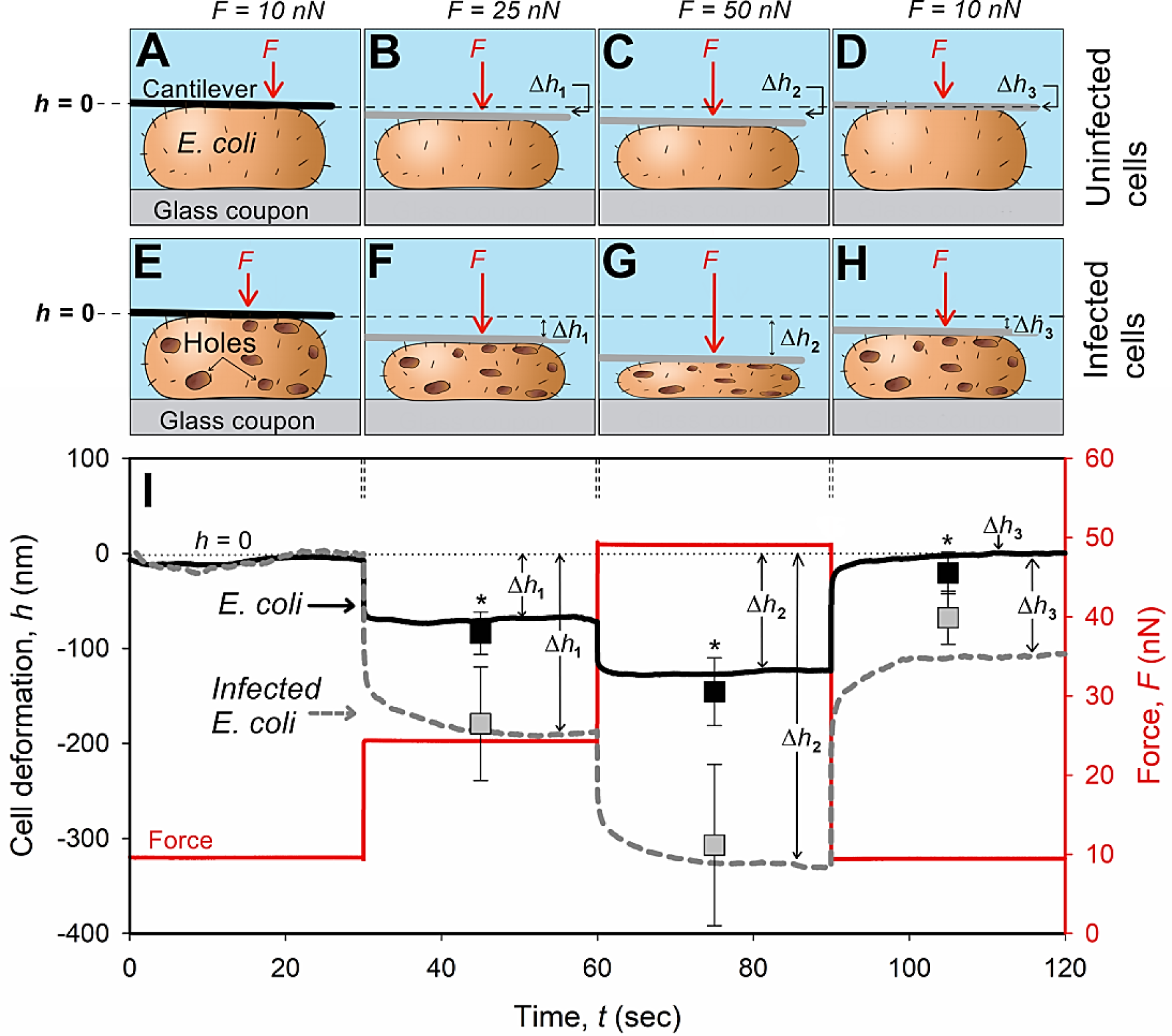
AFM measurements of applied force, *F*, and cell deformation, *h*, vs. time of applied force for infected and uninfected (carried out in separate experiments with no addition of T4 phages) *E. coli* cells. Two hours after the addition of T4 phages, infected *E. coli* were identified and analyzed according to the red fluorescence (*i*.*e*., “dead”) of cells with defined structure. (A-D) Schematic illustration of the tip-less cantilever as it deforms the uninfected and (E-H) infected *E. coli* cells at different applied forces. Black full lines (A, E) indicate the initial position of the tip-less AFM cantilever, while the gray lines (B-D, F-H) mark the position of that cantilever following the application of a specific force. (I) A representative curve where the deformation (indentation), *h*, was set to zero when the AFM cantilever applied *F*=10 nN. The deformations of uninfected or infected *E. coli* cells (Δ*h*_1_, Δ*h*_2_ and Δ*h*_3_) were measured by applying 25 nN, 50 nN, and 10 nN, respectively. Scatter plot indicate the average and standard deviation from 12 different bacteria measured at 3 to 5 independent experiments. Asterisks indicate a statistically significant difference between uninfected and infected *E. coli* cells (*p* < 0.01, *n*=12).

In addition to cell envelope stiffness, the elasticity (E) of a whole cell was calculated based on a strain response (deformation) of a single bacterium to a sudden change in force. These measurements were done *in-situ* using a tip-less AFM cantilever (Figs. 5, S3, S4), and the equation that was used to calculate *E* is specified in the Method section. Infected bacteria were identified two hours after the addition of T4 according to the red fluorescence of the cells (*i*.*e*., “dead” but intact bacteria). Compatible force profiles were also measured from uninfected (no addition of T4) bacteria two hours after placing the attached *E. coli* in the AFM bio-cell.

Initial contact between the tip-less cantilever and the bacteria was achieved by applying a constant force of 10 nN for 30 sec (Fig. 5I). During that time, the deformation of infected and uninfected bacteria was defined as *h*=0 nm (Fig. 5 A, E). A sudden (< 1 ms) change in force was then applied (from 10 nN to 25 nN), and F maintained constant for 30 sec. For uninfected bacteria, this resulted in an initial cell deformation, characterized by a stepwise decrease in cell height, of Δ*h*_1_ = 84±22 nm (Fig. 5B, I). Under the same applied change in force, infected bacteria deformed twice as much (Δ*h*_1_, 179±60 nm). Note, infected bacteria also deformed with an initial swift deformation followed by a slow (15-20 sec) and gradual decay (Fig. 5F, I), which is an indication of a viscous component that was not observed in uninfected bacteria. Concomitantly, the Young modulus (elasticity) of the infected cells was significantly (*p*<0.01, *n*= 15) lower (46%) than uninfected bacteria (33±13 kPa and 61±32 kPa, respectively). Increasing the applied force to 50 nN (additional step of 25 nN) resulted in a similar trend (Fig 5C, G); namely, greater (105%) deformation of infected cells compared to the uninfected bacteria (Δ*h*_2_, 307±85 nm and 146±36 nm respectively). Similarly to the above, the calculated *E* of infected cells was also significantly (P<0.01, n= 15) lower (41%) than uninfected bacteria (72 ± 20 kPa and 122 ± 54 kPa, respectively).

Retracting the cantilever to the initial force of contact (10 nN) caused uninfected bacteria to completely recover to their initial thickness (Δ*h*_3_, 21±22 nm) (Fig. 5D). In contrast, infected bacteria (with a define structure, Fig. 5H) remained significantly thinner after relaxing the applied force (Δ*h*_3_, 68±28 nm). Moreover, calculated elasticity following the recovery segment (Fig. 5I, 120 s > *t* > 90 s) remained much lower (48%) for infected cells than uninfected bacteria (56±18 kPa and 107±68 kPa, respectively).

## DISCUSSION

Studies of bacterial cell topography following phage infection have typically been carried at a nano-scale resolution by electron-based microscopy methods that are not suited to real time imaging ^18–21^. During the last decade, cell topography after phage infection was imaged by AFM-based techniques under dehydrated ^12,22^ and hydrated (*in situ*) conditions ^13,23,24^. In addition to nano-scale imaging, AFM has been used to measure changes in biomechanical properties of phage-infected bacteria ^13,23^. However, a major challenge of using *in situ* AFM to study bacterial samples is attaching the cells to the surface ^25–27^. One option is using cross-linking approaches (*e*.*g*., EDS-NHS and polydopamine) that immobilize the cells to the surface ^28,29^. However, these approaches might affect the physiological characteristics of biological samples. A common approach for biological samples is using a positively charged poly-L-lysine “bio-glue” to attach negatively charged cells with the surface. Unfortunately, it is challenging to control the thickness of the poly-L-lysine layer, which can be several µm thick, thus act as a soft (cushion-like) support that affects the elasticity measurements of the cells.

The cushion effect of poly-L-lysine was avoided here by using a rigid (vertical elasticity of ∼10 MPa) ^30^, positively charged LBL to act as a nano-thin glue (Fig. 1). The LBL adheres to *E. coli* cells via electrostatic interactions between the negatively charged lipopolysaccharides ^31^ and the positively charged LBL ^32^. We note that during imaging and elasticity measurements, the AFM tip exerts vertical and lateral forces on the cells. These forces can move and/or dislodge the cells from the surface, thus impairing the AFM measurements. However, the attachment of *E. coli* cells to the LBL layer under physiological conditions was robust enough to prevent cell motion and dissociation from the surface (Fig. 1), without impacting cell viability (Fig. 2A, 3A, S1A-B).

The earliest infection-specific changes (*t*<10 min) was the increase in SYTO 9 fluorescence, presumably reflecting the accumulation of newly synthesized phage DNA. This is consistent with previous studies, where, although host DNA is degraded to nucleotides during the early stage of infection, viral DNA biosynthesis is so great that total DNA mass, accumulates to levels ∼10-fold higher than pre-infection ^33^. Concurrently, a significant reduction in cell envelope stiffness (i.e., outer and inner membranes as well as the peptidoglycan layer) was measured (Fig. 4). This result was unexpected, since the generally accepted view is that the infected cell envelope is not challenged by the lysis proteins until the end of the infection cycle (i.e., before viron release). However, the infection event itself does result in damage to the host envelope; at each infection site, the outer membrane is penetrated by the puncturing device composed of a trimer of gp*5* attached at the end of the tail tube ^34–36^. Exact quantification of the multiplicity of infection (MOI) under these conditions is challenging, but experiments using similar physiological conditions suggest that only a few phages were adsorbed per cell (Fig. 1, *t*=20), far fewer than the high MOI needed to induce lytic events from without ^36,37^. Three gp*5* lysozyme domains are liberated in the periplasm per infecting phage; critical damage could be done to the peptidoglycan layer just from these molecules alone. Moreover, the penetration of the tail tube through the outer membrane may induce host envelope maintenance systems that could temporarily alter the physical properties of the cell. In addition, throughout the morphogenesis phase, the spanin complexes connecting the inner and outer membrane accumulate in the envelope ^10^. Although the membrane-fusion activity of spanins is blocked until the peptidoglycan layer is perforated after holin triggering, both holin and spanin comprise a direct protein-mediated link between the membranes. These links could affect the stiffness and elasticity of the host cell envelope. Finally, about half of the T4 genes have unknown function and could be participating in these early effects on the envelope properties. For example, phage lambda has three “moron” loci that modify the envelope early in infection ^38–40^. The powerful genetics available for experiments with T4 combined with Bio-AFM imaging should enable us to further examine the relationship between the expression and function of these genes in the early stages of *E. coli* infection.

One to two hours after the addition of T4, structural damages to the cell envelope were detectable by the penetration of PI (fluoresced in red) into the cells (*t*=60 to 120 min, Fig. 2, S1D). Taken together with the progression of the observed cell population from “live” to “dead” staining over a reasonable period (∼60 min), this result adds confidence that a robust phage infection was occurring in the infected cell population. Concurrently, numerous nanoscale ruptures (100-300 nm in diameter) on the outer cell envelope (15-40 nm in depth) and through the cell envelope (< 300 nm deep) were captured by AFM (Fig. 2H and I). These changes in cell topography are likely the result of T4 infection as no such damages were detected in uninfected cells (Fig 2A). Previously, comparable lesions (∼350 nm) were found in the inner membrane of *E. coli* cells after triggering of the T4 holin ^18,41^. These inner-membrane holes were found to be formed by T4 holin (gp*t*) at a programmed time, allowing the T4 endolysin to attack the peptidoglycan layer and control the timing of *E. coli* lysis. In turn, the destruction of the peptidoglycan layer liberates the spanin complexes, allowing them to diffuse laterally, aggregate and disrupt the structure of the outer membrane by fusion with the inner membrane ^9^. This pathway results in explosive, localized lysis in planktonic cells, with most “blow out” events localized to the cell poles in the case of phage lambda ^10^. The simplest interpretation of the results of the nano-topographical studies is that in a biofilm state, the outer membrane disruptions occur at random locales around the cell, possibly reflecting the underlying holin lesions in the inner membrane. Whether or not, these new findings confer an advantage for more effective dispersal of phage progeny from cells in mature biofilms remains to be seen.

Complementary to the nano-topographical analysis, force measurements indicated dramatic differences in cell elasticity between infected and uninfected bacteria (Fig. 5). Based on numerical fitting of the data to the classical models, such as Kelvin-Voigt, calculation of the host elasticity and viscosity yielded large errors for the viscous component(s) due to low sensitivity of the fitting method to changes in viscosity (Fig. 5I). Therefore, we considered only the elasticity of the *E. coli* cell. It should be noted, however, that in the case of viscoelastic materials (not plastic), viscosity only affects the time at which the strain reaches a steady state, while elasticity determines the strain at equilibrium. Therefore, to calculate the elasticity, E, we considered only the initial and final strains (30 sec after the sudden change in force). Nonetheless, slower deformation of infected cells under constant forces of 25 nN or 50 nN (Fig. 5I) indicated that infected cells exhibited viscoelastic or even plastic behaviour (i.e., cells deformation did not recover after relaxing the stress). On the other hand, uninfected bacteria exhibited pure elastic behaviour. We propose that the viscoelastic and plastic behaviour of infected cells is due to the production and assembly of new viral DNA during this infection phase ^42^. We surmise that it is likely that these infected cells remained partly deformed following the gradual loss of cytoplasm via lesions in the cell envelope. Yet, these infected bacteria did not explode during virion release, as the cells were not completely deformed, but instead partly flexed back after relaxing the applied force.

## CONCLUSIONS

The results of this study suggest that the lysis paradigm of T4, which was commonly studied based on planktonic cells, is markedly different for *E. coli* cells that are in the process of forming a biofilm. Planktonic *E. coli* cells lyse in an explosion, after which all the virions are instantly and simultaneously released from the cells. On the other hand, irreversibly attached *E. coli* cells, which are forming a biofilm, undergo a gradual (minutes to hours) lysis pathway. More importantly, during the lytic cycle of cells that are forming biofilm, the cell envelope becomes highly perforated and softer. Specifically, the elasticity of the cells decreases over time, and the cells exhibit more viscous and plastic (unrecovered deformation) behaviour than uninfected cells.

The explosive mechanism may provide ecological advantage for planktonic *E. coli* bacteria as the probability of phage progeny to re-infect neighbouring cells increases significantly. However, a controlled and gradual release may minimize the risk of lytic events from without for biofilm forming *E. coli* as virions are released in close proximity to other attached cells and diffusion is into the bulk is limited. Nonetheless, the clear differences in the lytic cycle of biofilm forming *E. coli*, including changes in the structural and biochemical properties of the cells pose new knowledge gaps, related to: (i) the impact of depolymerases expression in phage infection of multi-layered biofilms, (ii) the role of endolysins, holins and spanins in the lysis of bacterial biofilms and (iii) the effect of phage infection on the mechanical properties of a biofilms structure. We expect that elucidating the lysis phenomenon in bacterial biofilms may provide new means to enhance the efficiency of biofilm removal and phage therapeutic.

## METHODS

### Preparation of a lipid bilayer (LBL) surface to immobilize bacterial cells

AFM glass coupons (25 mm in diameter, Menzel Gläser, MENZCB00250RAC) were thoroughly cleaned with acetone and ethanol followed by Milli-Q water. The AFM coupons were then dried with N_2_ gas and UV cleaned for 10 minutes prior to the vesicle fusion on the glass substrate. Zwitterionic 1,2-dimyristoyl-sn-glycero-3-phosphocholine (DMPC), and positively charged 1,2-dimyristoyl-3-trimethylammonium-propane (DMTAP) lipids (Avanti Polar Lipids, Inc., Alabama, USA) were used for the LBL coating. NaCl solution (1 ml, 150 mM) was added to the dried lipid combination (DMPC + DMTAP), resulting a positive surface charge ^32^, to reach a total concentration of 0.5 mM lipids. Ultra-sonication for 2 min (60 Hz; Witeg-Germany), followed by gradual heating (55 °C) for 20 min, was performed to achieve a well-dispersed vesicle solution. The AFM glass coupon was uniformly covered by the vesicle solution (350 µl) and incubated at room temperature for 10 minutes. During that time vesicle were fused with the surface, leading to a self-assembled LBL. The residual vesicles were removed by washing with 150 mM NaCl solution.

### Bacteria culture and AFM bio-cell setup

*E. coli* B strain (ATCC 11303) was cultivated to exponential phase (OD_600_ of 0.6-0.8) in Luria-Bertani broth (BD244620; Becton, Dickinson and Company, United States) at 37 °C. *E. coli* cells were irreversibly attached by placing a subsample (∼600 µL) onto an LBL pre-coated AFM glass coupon (Fig. 1). Inoculated AFM coupons were kept in a humid chamber (to minimize cell dehydration) at 37 °C for ∼8 h. At the end of the incubation any loosely bounded bacteria were washed with sterile phosphate buffer saline PBS, P4417, Sigma, US). Immobilized bacteria were stained with SYTO 9 (final concentration of 0.01 mM, Invitrogen, ThermoFisher Scientific, United States), a fluorescent fluorophore (Ex 485 nm, Em 498 nm), for 20 min in the dark. Excess stain was gently removed by replacing the PBS media three times.

The glass coupon with immobilized and stained bacteria was assembled into an AFM bio-cell according to manufacture instructions (JPK Instruments, Germany) and filled with sterile PBS (1 ml) (Fig. S5). Bacteria were initially localized by an inverted epifluorescence microscope (Zeiss Axio Observer.Z1, Germany) equipped with a green filter (Ex. 470±20 nm and Em. 525±25 nm) and integrated with the AFM. Nano-scale structures and mechanical properties of localized bacteria were captured and analyzed *in-situ* and in real time by an AFM for ∼4 h (JPK Instruments AG, Berlin, Germany). SYTO 9 (Ex. 485 nm and Em. 498 nm) and PI (Ex. 535 nm and Em. 617 nm), (Live/Dead BacLight Bacterial Viability and Counting Kit, Invitrogen, ThermoFisher Scientific, United States) were added to the bio-cell at concentration of 0.06 mM (Fig. S5). The samples were stained for 20 min in the dark. Excess stain was removed by replacing the PBS media three times. Images of Live/Dead cells were captured by epifluorescent microscopy, while nano-scale images and force measurements were captured and analyzed by AFM at different intervals throughout the experiment (between 10 to 240 min). Detailed information on AFM imaging and measurements are provided below.

### Calculation of the cell elasticity using a tip-less cantilever

The elasticity (*E*) was calculated by:

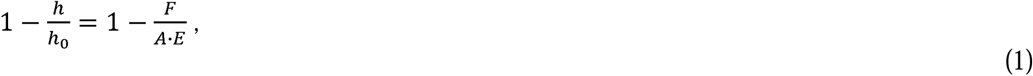

where h0 is the cell height (thickness) as measured by AFM imaging and h is the indentation depth of the tip-less cantilever into the cell (see Fig. S4). F is the force that was applied on the cell, as measured by the AFM. A is the contact area between the tip-less cantilever and the cell. Note, A changes with h according to Eq. S2. Additional details on the elasticity calculations are provided in Section 4.3 in the Supporting Information.

### Propagation of T4 phages and proceeding of the infection experiments

T4 phages (ATCC, 11303-B4) were thawed and propagated according to Adams et al. ^43^. Briefly, *E. coli* (1 ml, log phase) were mixed with 50 µL T4 phage and 5 ml worm soft agar (30 mg/ml of tryptic soy broth (TSB) + 0.5 % agar), pour onto TSA plates and incubated at 37°C overnight. Sterile PBS (10 ml) was added into the inoculated LB plate for 1h at room temperature. PBS and phages (∼8ml) were collected and centrifuged (8000 × g) for 15 minutes. The concentrated phages were filtered through a 0.45 µm filter (SLHVM33RS, Millex-HV Syringe Filter Unit, Merck, Germany) and stored at 4°C.

Phage concentrations in the stock culture were quantified by counting plaque forming units (PFU). Briefly, log phase *E. coli* (1 ml) was mixed with a serial dilution of T4 phages in 5 ml of warm 0.5% TSB agar. The mixed media was poured onto a solid TSB (1%) plate and incubated at 37°C overnight. Plaques were counted after 12h. Negative controls (*i*.*e*., no addition of T4) were done for all PFU test.

Following the initial *E. coli* localization and characterization, T4 phage were added (3×10^8^ phages ml^-1^) to the AFM bio-cell together with SYTO 9 and PI (BacLight Bacterial Viability and Counting Kit, Invitrogen, ThermoFisher Scientific, United States) at concentration of 0.06 mM. Images of infected cells were first captured by epifluorescent microscopy followed by nano-scale images and force measurements of the same cells were captured in throughout the experiment. Transmission electron microscopy micrographs were also captured in similar time intervals.

### AFM imaging and force *vs*. distance *vs*. time measurements

AFM images were acquired using JPKSPM (JPK Instruments AG, Berlin, Germany) with an SNL-10 probe (Bruker, Camarillo, CA) and a spring constant of 0.35 N/m in quantitative imaging (QI) mode at room temperature (25° C). The AFM force *vs*. distance *vs*. time curves of entire cells were measured using a tipless cantilever (MIKROMASCH, Sofia, Bulgaria). The spring constant of the cantilever used here is 0.4 N/m. The deflection *vs*. displacement *vs*. time data were converted into force *vs*. relative distance *vs*. time graphs using the JPKSPM-data processing software. Fluorescent images of the samples on the substrates were captured by an epifluorescent microscope (Axio Zoom. V16, Zeiss, Germany) which is coupled with the AFM.

### Transmission electron microscopy (TEM)

*E. coli* B cells (600 µl) were grown for 8 h on 25 mm, 0.2 µm Supor® PES membrane disc filters (Pall Corporation, USA), that were placed in a petri-dish in a humid chamber at 37°C. Loosely attached cells were removed from the membrane by rigors washing with a 10 ml syringe field with sterile PBS. Immobilized *E. coli* cells were inoculated with T4 (final concentration of 5×10^7^ phages ml^-1^). At t=0 before phage infection and at 20 min intervals after infection, a filter was fixed with glutaraldehyde solution (2.5%; G7651, Sigma-Aldrich, United States) in a 1.5 ml eppendorf overnight at 4 °C.. Fixed samples were stained with osmium tetroxide and sequentially dehydrated according to Bar-Zeev et al. 2015 (38). Dried samples were embedded in an epoxy resin (Embed 812, Electron Microscopy Sciences, Hatfield, PA), cut into 70 nm slices and imaged with a Philips CM12 TEM equipped with a Gatan 791 CCD camera.

### Statistical analysis

Statistical analyses were performed using Microsoft Excel and the data were plotted using Origin 2018 or by SigmaPlot. Standard deviation, averaging and T-test were conducted by this excel sheet. Independent bacterial samples were measured to determine cell deformation (n=12), cell stiffness (n=12), and elasticity (n=15). Further, confidence level of p < 0.01 was required for the T-test analyses throughout the manuscript.

## ACKNOWLEDGMENT

We would like to thank the Roskind fund and the Roy J Zuckerberg Career Development fund for their financial assistance. The authors gratefully acknowledge Dr. David Lowry and the Electron Microscopy division of the CLAS Bioimaging Facility at ASU. Ry Young thanks the National Institute of General Medical Science, Grant # R35GM136396-01.

## Author Contributions

Conceived and designed the experiments: E. B-Z., Y.K. and S.A. Performed the samplings: E. B-Z., Y.K., and S.A. Analyzed the data: E. B-Z., Y.K., F.P., R.Y., and S.A. Contributed reagents/ materials/analysis tools: E. B-Z., Y.K. F.P. and S.A. Wrote the paper: E. B-Z., Y.K., F.P., R.Y., and S.A.

## Competing Interests Statement

The authors declare no competing financial and non-financial interests.

## Data availability

All data measured and analyzed during this study are included in the manuscript.

## Notes

### Competing Interest Statement

The authors have declared no competing interest.

