## Supporting Information for "Bursting out: linking changes in nano-topography and biomechanical properties of biofilm-forming Escherichia coli to T4 lytic cycle"

*eLife*

**Shiju Abraham[1], Yair Kaufman[1*], François Perreault[4], Ry Young[2,3], and Edo Bar-Zeev[1*]**

[1]Zuckerberg Institute for Water Research, The Blaustein Institutes for Desert Research, Ben-Gurion University of the Negev, Sde Boqer Campus, Midreshet Ben-Gurion, 8499000, Israel.
[2]Department of Biochemistry and Biophysics, Texas A&M University, College Station, TX, 77843, USA
[3]Center for Phage Technology, Department of Biochemistry and Biophysics, Texas A&M University, College Station, TX, 77843, USA
[4]School of Sustainable Engineering and the Built Environment, Arizona State University, Tempe, AZ 85287-3005, United States

### RESULTS AND DISCUSSION

**1. Control experiments:** Our results indicate that the green florescence ("live") of attached *E. coli* cells to the positively charged lipid bilayer (LBL) was mostly unchanged during 120 min (Fig. S1 A,B panels). However, a few bacteria were found to become red ("dead") over time, indicating that some cells were physiologically impaired. On the other hand, *E. coli* cells showed an increase in green florescence following 10 to 60 min after the addition of T4 phages (Fig S1 C), which was likely due to the accumulation of intracellular T4. In addition, many cells became red ("dead") following 60 to 120 min after T4 addition, indicating cell wall perforation (Fig S1 D).

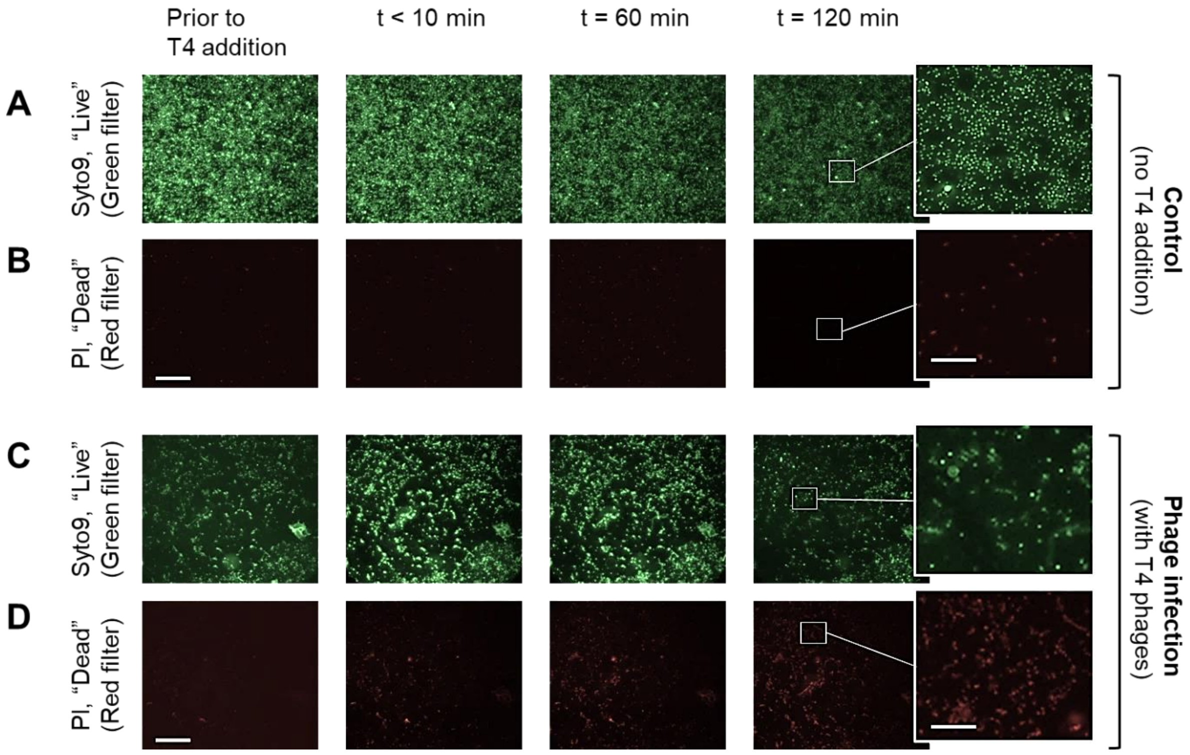


**Figure S1.** Live (green) and dead (red) staining of *E. coli* cells that were attached to the positively charged LBL and captured *in-situ* by epifluorescence microscopy. (A and B) Control experiments were carried without the addition of T4 phages for ~120 min in the AFM bio-cell. (C and D) Concomitant experiments included incubation of attached cells with T4 phages for 120 min. Scale bar of the main plots is 50 µm and of the inserts is 20 µm.

Nano-scale images of *E. coli* cells were captured over time without the addition of T4 phages to test the effect of the positively charged LBL on cell morphology. Our images indicate that nano-scale morphology remained unchanged over time. Hence, we surmise that attaching the cells to the coated LBL layer did not affect the morphology of the cells.

**2. Cell dimensions of uninfected *E. coli* bacteria.** Overall, uninfected *E. coli* cells were found to be around 1 µm in diameter and a length that ranged from 2 µm to 3 µm (Fig. S2).

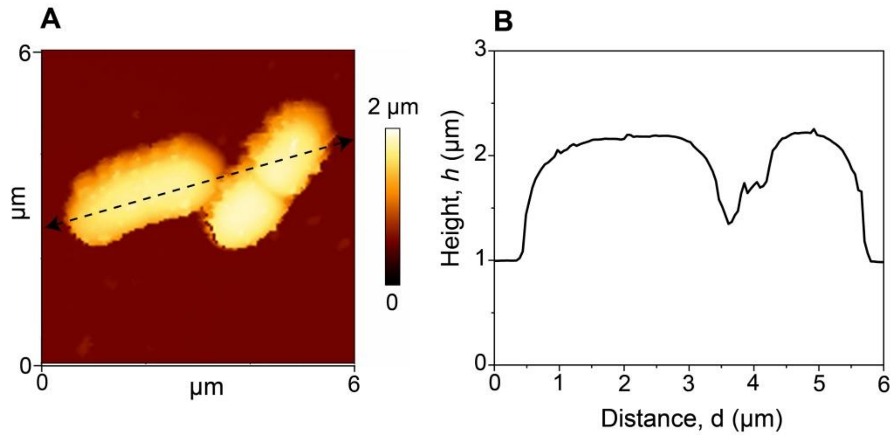


**Figure S2.** (A) AFM topography image of uninfected *E. coli* cells. (B) Height profile (cell cross-section) corresponding to the black dotted line in (A). Similar images were captured for numerous bacteria.

### 4. METHODS

***4.3 Calculation of the cell elasticity of a single E. coli cell.*** To calculate the elasticity of a *E. coli* cell, $E$, the cell was modeled as a cylinder, as depicted in Fig. S3 A.

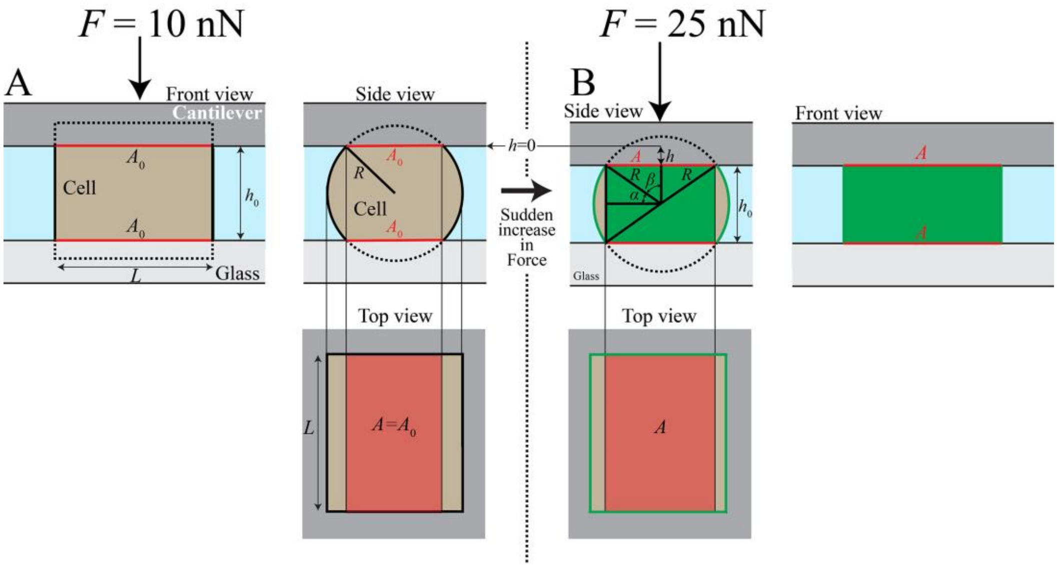


**Figure S3**. Using a tip-less AFM cantilever, a sudden change in force was applied on a separated *E. coli* cell to measure the elasticity, $E$. (A) The cell was modeled as a cylinder with a base radius $R$, and length, $L$. As the cantilever applied force on the cell, as a first order approximation, it was assumed that the parts of the cell that was not in contact with the cantilever, was not deformed; however, the contact area with the cantilever, $A$ (red), increases with $F$, i.e., the applied pressure on the cell decreased with the cantilever indentation, $h$.

The strain response of a single cell to a sudden change in force was modeled using Kelvin-Voigt in addition to water permeability through the cell membrane:

$$1-\frac{h}{h_0}=1-\frac{F}{A\ \cdot E} \tag{S1}$$

where $h_0$ is the height of the cell measured by AFM, $h$ is the indentation of the cantilever into the cell (Fig. S3 B), $F$ is the force that was applied by the AFM cantilever on the cell, and $A$ is the cantilever-cell contact area that increases with $h$ (red in Fig. S3).

Based on the simplified model in Fig. S3, $A$ was estimated using the following equation:

$$A=A_0+L\sqrt{2Rh-h^2}, \tag{S2}$$

where $A_0$ (estimated parameter) is the cantilever-cell contact area before the sudden change in $F$ (at $t=0$), the angles $\alpha$ and $\beta$ are depicted in Fig. S3 B and they are given by $\sin\alpha=\left(\frac{h_0}{2}-h\right)/R$ and $\cos\beta=\frac{R-h}{R}$, $R$ is the radius of the cylinder base, and $L$ is the cell length (Fig. S3 A and B).

Fig. S4 shows typical deformation/strain measurements of non-infected *E. coli* cell and after an hour and a half (~90 min) since the addition of T4 to the *E. coli* cell culture (infected cell). The red circles are the calculated deformations for the two cells at *t*=0 and at *t*=30 s.

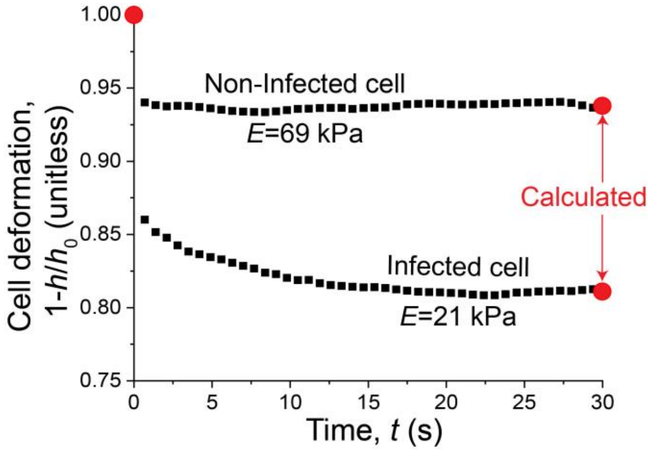


**Figure S4:** Deformation/stain response of *E. coli* cells to a sudden change in stress/force. The black rectangles are the measured data, and the red circles were calculated using Eq. S1 at *t*=0 and at *t*=30 s.

***4.1 Imaging the physiological state of E. coli cells using following T4 phages addition.*** The Live/Dead kit (Invitrogen, ThermoFisher Scientific, United States) is commonly used to estimate the physiological state of bacterial cells. Briefly, the kit is based on two molecules with specific florescence spectra and different penetration efficacy into the cell. Syto9 is a small, fluorescently green (Ex. 485 nm and Em. 498 nm) molecule that penetrates all cells (including those with intact cell-wall). Differently, Propidium iodide (PI) florescence in red (Ex. 535 nm and Em. 617 nm) and is relatively larger than Syto 9, thus penetrate the bacteria only if the cell-wall was perforated. Hence, live cells will appear to be green, while dead cells (or with impair cell-wall) will appear red (Fig. S1). Additional information is provided by the Invitrogen (USA) for L7007 Live/Dead® BacLight Bacterial Viability Kit).

***4.2 Experimental set up for the attachment of E. coli cells on lipid bilayer (LBL) surface.*** *E. coli* bacteria must be immobilized to an AFM glass coupon to visualize cells and while conducting mechanical measurements in real time and *in-situ* (namely in phosphate-buffered saline, PBS) using AFM. Since the AFM glass coupon and the surface of the bacterial cells are often negatively charged, it is common to use a support layer which is positively charged to irreversibly attach the cells. However, it is highly important that the coated support layer will not cause physiological stress to the attached cells. Further, reducing the effect of the support layer on the overall elasticity and thickness of attached bacteria is critical to gain equate force measurements. Here, we adopted a novel approach to immobilize bacteria on the surface by employing a net positive surface charge lipid bilayer, LBL (Fig. 1). Bellow we provide a schematic overview of the steps applied before visualizing and analyzing the samples by AFM (Fig. S5). Coating the glass AFM coupon by LBL requires that the surface will be thoroughly cleaned (Fig. S5 A). Thus the glass surface was washed with acetone, ethanol and Milli-Q water, dried with $N_2$ gas followed by UV treatment for 10 min. The clean AFM coupon was then coated by a positively charged LBL, additional information is provided in the main text (Fig. S5 B). *E. coli* cells (at an exponential phase) were deposited on the surface for 8 hours. The AFM coupon was then intensively washed to remove all the cells that were loosely attached (Fig. S5 C). AFM coupon with the attached cells was then mounted into the AFM biocell (Fig. S5 D) before the initiation of the experiments.

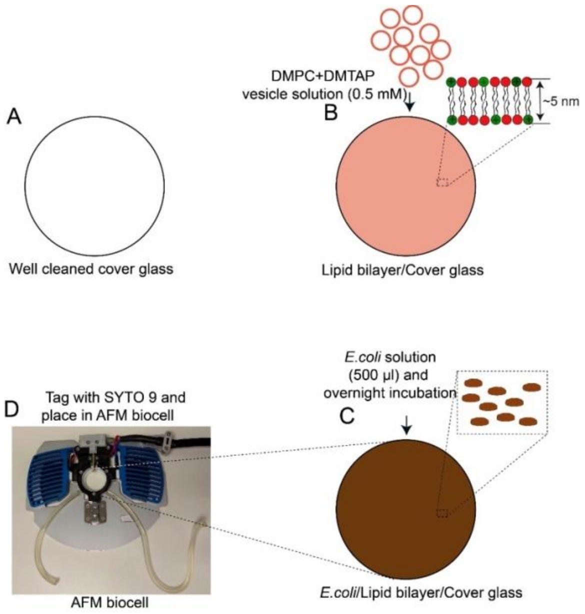


**Figure S5**. Schematic illustration of the experimental setup used to attach *E. coli on* an AFM glass coupon. The main steps included: (A) thorough cleaning of an AFM glass coupon; (B) formation of a positively surface charged LBL on the AFM glass coupon; (C) attachment of *E. coli* bacteria to the LBL substrate; (D) Assemble the coupon with the attached cells into the biocell before visualization and force measurements.
